# A refined sticker-spacer grammar marks disordered regions with interpretable pathogenic missense variation

**DOI:** 10.64898/2026.09.25.754449

**Authors:** Gavin E. Remo, Gabriel A. Andaverde, Karen A. Lewis, Loren E. Hough, Steven T. Whitten

## Abstract

Intrinsically disordered regions (IDRs) of proteins show low sequence conservation, and sequence position alone is a weak predictor of disease risk. Phase-separating IDRs (PS-IDRs) are an exception, showing nearly three times higher pathogenic mutation rates than typical IDRs. Here we determine which sequences within that class show the greatest enrichment of pathogenic missense variants and what local features they share. Using pathogenic mutation rates as an independent biological readout, we isolated a core PS-IDR subclass that maximizes pathogenic enrichment. These sequences are defined less by bulk amino acid content than by a refined sticker-spacer grammar in which aromatic residues are embedded in short glycine-rich motifs, while homotypic charge tracts are avoided. Motifs such as YGG/YGGG, FGG/FGGG, and related patterns are elevated 7-to >200-fold relative to conventional IDRs. Comparison of pathogenic and benign substitutions then shows why genetic variation in IDRs has been confounding: the grammar itself is almost never mutated. Instead, the principal pathogenic targets are glycines directly flanked by non-glycine residues. These findings link a specific local sequence architecture to mutational vulnerability and provide a practical framework for interpreting variants in disordered proteins.

**SIGNIFICANCE:** Disease-causing mutations in unstructured protein regions are common, yet hard to interpret because those sequences look loosely constrained. This study shows that a subset of condensate-forming regions follows a precise molecular grammar. Aromatic residues sit in short glycine-rich motifs that are almost never mutated, while isolated glycines are the main pathogenic targets. That distinction identifies where pathogenic missense variants concentrate in disordered proteins.

## INTRODUCTION

Missense mutations in intrinsically disordered regions (IDRs) are common in the human proteome, yet their functional consequences remain difficult to interpret (1–3). IDR sequences are, on average, less conserved than those in folded domains and therefore experience higher levels of genetic variability (4, 5). Even so, certain disordered segments harbor a disproportionately high number of pathogenic mutations. For example, missense mutations in TDP-43 and FUS, RNA-binding proteins with prion-like low-complexity domains, are strongly associated with amyotrophic lateral sclerosis and frontotemporal dementia (6–10). This pattern indicates that specific sequence features can render some IDRs particularly vulnerable to pathogenic substitutions. Defining such features is essential for understanding how genetic variation contributes to disease.

Protein liquid-liquid phase separation, or more generally protein phase separation, has emerged as a key functional mechanism for many IDRs (11, 12). Phase-separating IDRs (PS-IDRs) drive the formation of dynamic, membraneless condensates that organize cellular biochemistry and participate in diverse processes, including stress responses, ribosome biogenesis, chromatin regulation, RNA processing, and signal transduction (13–18). Dysregulation of phase separation has been linked to multiple human diseases, including cancer, neurodegeneration, and autism spectrum disorder (2, 19, 20). Consistent with this connection, disease-associated proteins show a higher prevalence of predicted phase separation behavior than is typical for the human proteome, and pathogenic missense mutation rates are higher in predicted PS-IDRs relative to other IDRs (21). PS-IDRs are therefore more mutationally vulnerable, on average, than non-phase-separating IDRs (nonPS-IDRs). It is not known whether this enrichment of pathogenic mutations is spread uniformly across all PS-IDRs or concentrated in a subset with distinctive local sequence features.

To distinguish those alternatives, we used pathogenic mutation rates as a biological filter. Sequence groups for folded regions, nonPS-IDRs, and PS-IDRs were varied systematically with ParSe version 2, a sliding-window classifier that uses hydrophobicity (*φ*) to separate folded from disordered segments and combines α-helix propensity with a sequence-based model of the polymer scaling exponent (*ν*_model_) to separate conventional ID from PS-ID (22). This strategy identified a narrow PS-IDR subclass that concentrates the pathogenic mutation signal. An independent, mutation-centric definition of functionally critical IDRs, requiring only that a pathogenic substitution be flanked by disordered sequence, mapped to the same region of property space. Two orthogonal amino acid property scales then recovered the high-pathogenicity subclass across the proteome. Within that subclass, aromatic residues were not randomly distributed but were instead preferentially embedded within glycine-rich contexts. This revealed a highly refined sticker-spacer grammar (23, 24), characterized by a strong enrichment of motifs such as Y, F, or W placed directly next to or inside G-runs. In contrast, homotypic charge tracts are avoided.

Comparison of pathogenic and benign missense mutations within these sequences further showed that the most enriched sticker-spacer motifs are rarely mutated. Isolated G, a glycine flanked on both sides by non-G residues, is the main site of pathogenic substitution. The motifs themselves are consistent with a local architecture that can remain disordered while still supporting condensate-forming interactions. More generally, the results illustrate how pathogenic mutation rates can resolve, inside disordered sequence, which local patterns are informative for variant interpretation.

## RESULTS

### A narrow PS-IDR subclass concentrates the pathogenic mutation signal

Previous studies have shown that pathogenic missense mutation rates are elevated in PS-IDRs relative to conventional IDRs (21). We hypothesized that these rates would increase further with a sequence’s computed PS potential, as higher potential reflects both greater predictive confidence and a narrower compositional window. To test this, we applied ParSe v2 (22, 25), our sequence-based algorithm for predicting domain-level protein organization for folded, ID, and PS-ID regions.

ParSe v2 employs a sliding-window approach (25 residues) across the primary sequence (Figure 1). Within each window, average *φ* (derived from globular protein contact matrices (26)) distinguishes folded (F) from intrinsically disordered (ID) segments. Intrinsic α-helix propensity (from native protein X-ray data (27)) combined with *ν*_model_ (based on hydrodynamic size (28)) then separates conventional ID regions (D) from those predicted to drive phase separation (P). Window labels are assigned to the central residue, with terminal residues receiving the labels of the flanking windows. Contiguous segments of ≥20 residues that are ≥90% one label are classified as folded, ID, or PS-ID domains.

**Figure 1.**
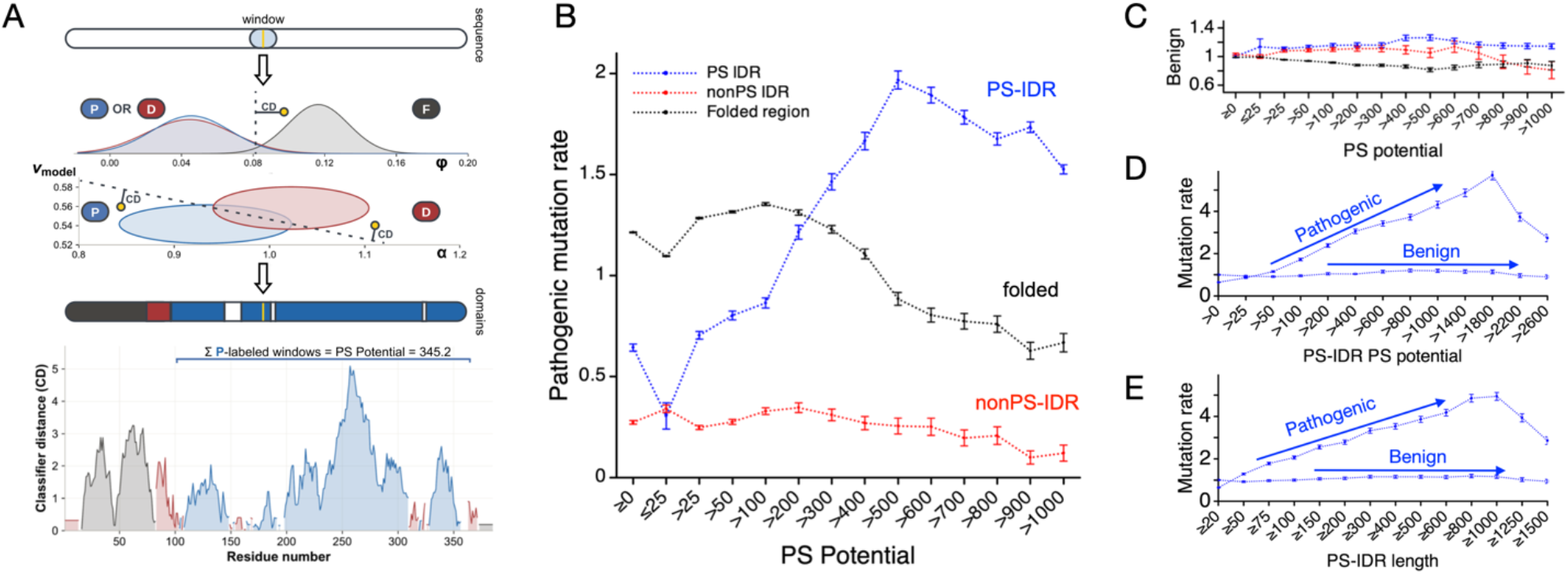
Domain labels from ParSe v2, and pathogenic missense rates as a function of PS potential. (**A**) ParSe v2 sliding-window analysis. For each contiguous 25-residue window, ParSe v2 calculates hydrophobicity (*φ*), α-helix propensity (*α*), and the polymer scaling exponent (*ν*_model_). Distributions for the folded (black), conventional ID (red), and PS-ID (blue) training sets are shown; decision boundaries are indicated by dashed lines. Windows are labeled folded (F), PS-ID (P), or conventional ID (D) according to their position relative to these boundaries, with the label assigned to the central residue. Contiguous segments ≥20 residues that are ≥90% one label define predicted domains. Classifier distance (CD) quantifies how far a window lies from its classification boundary. Summed CD values for P-labeled windows yield the sequence PS potential. (**B, C**) Normalized pathogenic (B) and benign (C) missense mutation rates by region class for proteins of varying overall PS potential. (**D, E**) Normalized pathogenic and benign missense mutation rates within PS-IDRs as a function of individual PS-IDR PS potential (D) or residue length (E). Error bars are the standard error.

The PS potential of a sequence is the summed classifier distance across all P-labeled windows. Classifier distance measures how far a window’s feature values lie from the relevant decision boundary used for its classification. Windows lying close to a boundary fall within the overlap region between training-set distributions and therefore carry greater uncertainty, whereas windows lying farther from the boundary have higher-confidence assignments. A higher “potential” can arise from longer predicted PS-IDRs, a greater number of predicted PS-IDRs, and higher confidence in these predictions. Proteins with PS potential ≥100 are identified as PS-predicted, consistent with the threshold observed for 43 experimentally validated homotypic phase-separating proteins (22). ParSe v2 shows good agreement with other ID (29, 30) (Figure S1) and phase-separation predictors (31–38) (Figure S2).

Confirmed pathogenic and benign missense mutations were obtained from the humsavar index of human genetic variations from UniProt (39). Normalized mutation rates were computed as the proportion of mutations in a region class divided by the proportion of residues in that class for a protein set. For the full reference human proteome, normalized pathogenic missense mutation rates were 0.26 ± 0.009 (IDRs), 0.65 ± 0.018 (PS-IDRs), and 1.21 ± 0.002 (folded regions; rate ± standard error). When proteins were binned by overall PS potential, rates within PS-IDRs rose sharply with increasing potential (Figure 1). For proteins with PS potential ≤25, the PS-IDR rate (0.28 ± 0.06) was comparable to conventional IDRs (0.33 ± 0.02). Rates within PS-IDRs increased dramatically for high-potential proteins, reaching 1.99 ± 0.04 (PS potential >500), exceeding even folded-region rates. Benign mutation rates showed little dependence on region class or PS potential. Similar trends held when binning by individual PS-IDR potential, rather than whole-sequence PS potential, or when considering domain length. Rates of pathogenic mutation within conventional IDRs remain largely unchanged across the stratification. Folded domains dominate the global mutational burden, consistent with structural constraints (40, 41). However, the strongest mutational vulnerability shifts dramatically to PS-IDRs in proteins enriched for phase separation behavior. Benign rates do not follow that shift. A general increase in mutability, or in the chance that a variant is simply observed, would raise pathogenic and benign labels together. The benign rate stays flat. The pathogenic enrichment is therefore better read as mutational vulnerability arising from functional constraint. Substitutions in these sequences are more likely to be costly, and that constraint is strongest in high-potential PS-IDRs.

### Mutational vulnerability is highest in hydrophilic PS-IDRs

As a second approach to identify the PS-IDR subsets with elevated mutational vulnerability, we systematically varied the ParSe v2 decision boundaries and tracked changes in region-specific pathogenic mutation rates (Figure 2). We first shifted the folded-versus-ID boundary (default *φ* = 0.083) in unit steps equal to the standard deviation of *φ* values in confirmed folded sequences (σ ≈ ±0.017). Negative shifts move the boundary to lower hydrophobicity; positive shifts move it higher. The combined pathogenic mutation rate across all IDRs (PS + conventional) reaches a minimum near the default cutoff. Shifts to higher *φ* raise that rate by mixing high-rate folded sequence into the ID pool, indicating that the default boundary avoids that contamination. Shifts to lower *φ* also raise the combined rate, but by a different mechanism. Pathogenic mutations concentrate in a hydrophilic PS-IDR subset. At every tested position, the PS-IDR pathogenic rate exceeds that of conventional IDRs, confirming that PS-IDRs form a functionally distinct subclass whose sequence properties correlate with elevated mutational vulnerability. Benign mutation rates remained essentially unchanged across all boundary positions (Figure S3). The pathogenic enrichment therefore tracks how the sequence classes are defined, not a change in background mutability.

**Figure 2.**
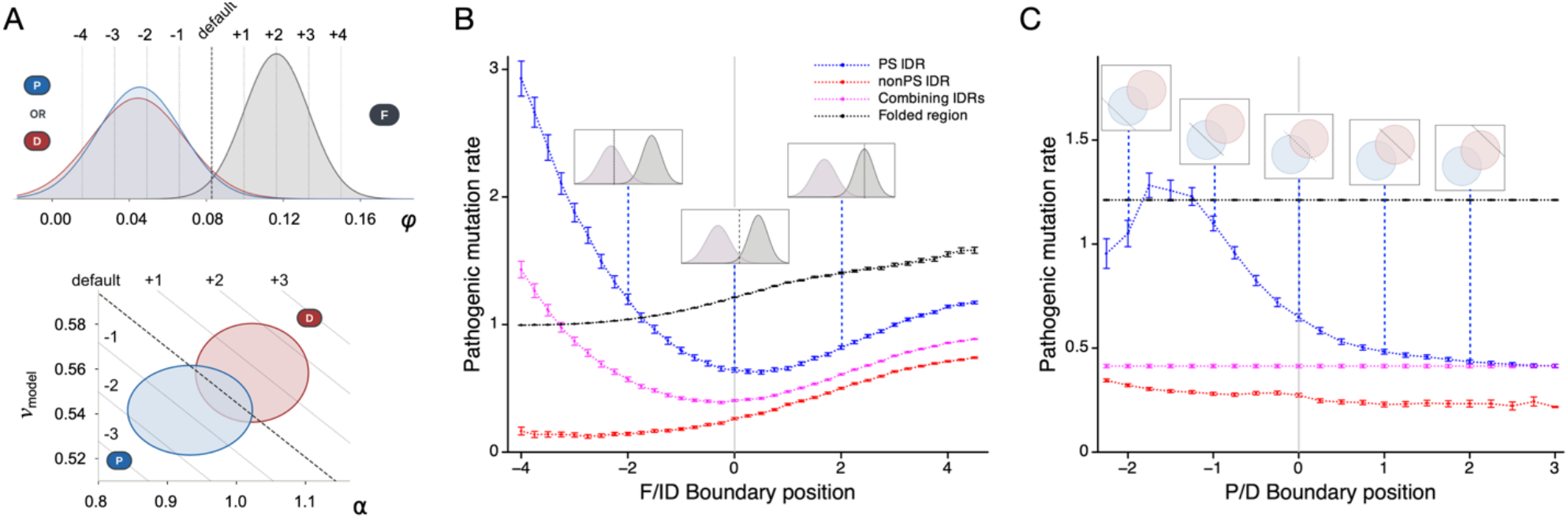
Pathogenic mutation rates are most strongly enriched in a narrow, compositionally selective subset of PS-IDRs. (**A**) Schematics of ParSe v2 decision boundary shifts. Top: folded-versus-ID (*φ*) boundary shifts, shown relative to the folded (black), PS-ID (blue), and ID (red) training set distributions. Bottom: P-versus-D boundary shifts in α versus *ν*_model_ property space, shown relative to the PS-ID (blue) and ID (red) training set centroids (mean ± standard deviation). (**B**) Normalized pathogenic mutation rates versus F/ID boundary position for PS-IDRs (blue), conventional IDRs (red), PS + conventional IDRs (cyan), and folded regions (black). (**C**) Normalized pathogenic mutation rates versus P/D boundary shifts for the same sets. Error bars are the standard error.

The same boundary shifts change the folded class in the expected direction. As the cutoff moves to higher *φ*, folded windows become progressively restricted to the most hydrophobic segments, and the folded pathogenic rate rises, consistent with disruption of tight side-chain packing in globular cores (40, 41). Extreme negative shifts make the cutoff highly permissive, eventually labeling nearly the entire proteome as folded and the normalized folded pathogenic rate approaches 1. Under these conditions, the remaining PS-IDRs are confined to the most extremely hydrophilic windows. Those sequences must remain disordered and soluble while still supporting the multivalent cohesive interactions that drive condensation. Their normalized pathogenic mutation rate (∼3) is among the highest seen in this study, implying that the need to stay disordered, soluble, and still able to cohere is mutationally vulnerable and functionally constrained.

Conventional IDRs showed only trivial changes in pathogenic rate across the full range of folded-versus-ID boundary shifts. Positive shifts (higher *φ* cutoff) mislabel some borderline-folded windows as disordered, modestly increasing the conventional IDR rate. Negative shifts shrink the disordered pool to only the most ultra-disordered, extremely hydrophilic windows, keeping the conventional IDR rate low and stable.

We next held the folded-versus-ID boundary fixed at its default value and shifted the internal P-versus-D boundary (slope preserved) in normalized units relative to the mean of confirmed PS-ID sequences in α-helix propensity and *ν*_model_. Negative shifts shrink the size of the PS-ID class; positive shifts enlarge it. The combined pathogenic rate across all IDRs remains flat (∼0.4), confirming that boundary movement merely repartitions the same residue pool. The PS-IDR pathogenic rate shows a clear asymmetric peak at moderate negative shifts, between -1 and -2, with enrichment falling sharply once the boundary passes the PS-ID training set centroid. Conventional IDRs exhibited only minimal changes in pathogenic rate with P-versus-D boundary shifts. Benign rates again remained largely unaffected (Figure S3).

Taken together, both boundary analyses show that the pathogenic signal is strongest when the PS-ID class is small and compositionally selective. Sequences deepest in PS-ID property space are among the most prone to pathogenic mutations in the human proteome, exceeding even folded domains.

### Disordered regions flanking pathogenic mutations define a distinct protein class

To further test whether the most disease-relevant IDRs are enriched for phase-separation properties without presupposing a ParSe domain classification, we identified all pathogenic missense mutations in humsavar that are flanked on each side by 25 disordered residues (≥90% P- or D-labeled). This procedure yielded 1,026 “pathogenic-site IDRs” (Figure 3). Of these, 733 (71%) were ≥90% P-labeled (PS-IDRs), whereas only 66 (6%) were ≥90% D-labeled (conventional IDRs). The remaining fraction consisted of mixed P/D segments. The local sequence properties of these pathogenic-site IDRs are shifted toward lower *φ* (more hydrophilic) and closely overlap the PS-ID training set distributions in α-helix propensity and *ν*_model_. Thus, an independent, mutation-centric definition of functionally critical IDRs converges on the same region of property space identified by the boundary-shifting analysis, indicating that the high-pathogenicity sequences are not an artifact of the ParSe classification itself. These results show that the sequences surrounding pathogenic missense mutation sites in disordered regions combine extreme hydrophilicity with the low α-helix propensity and polymer scaling behavior characteristic of PS-IDRs.

**Figure 3.**
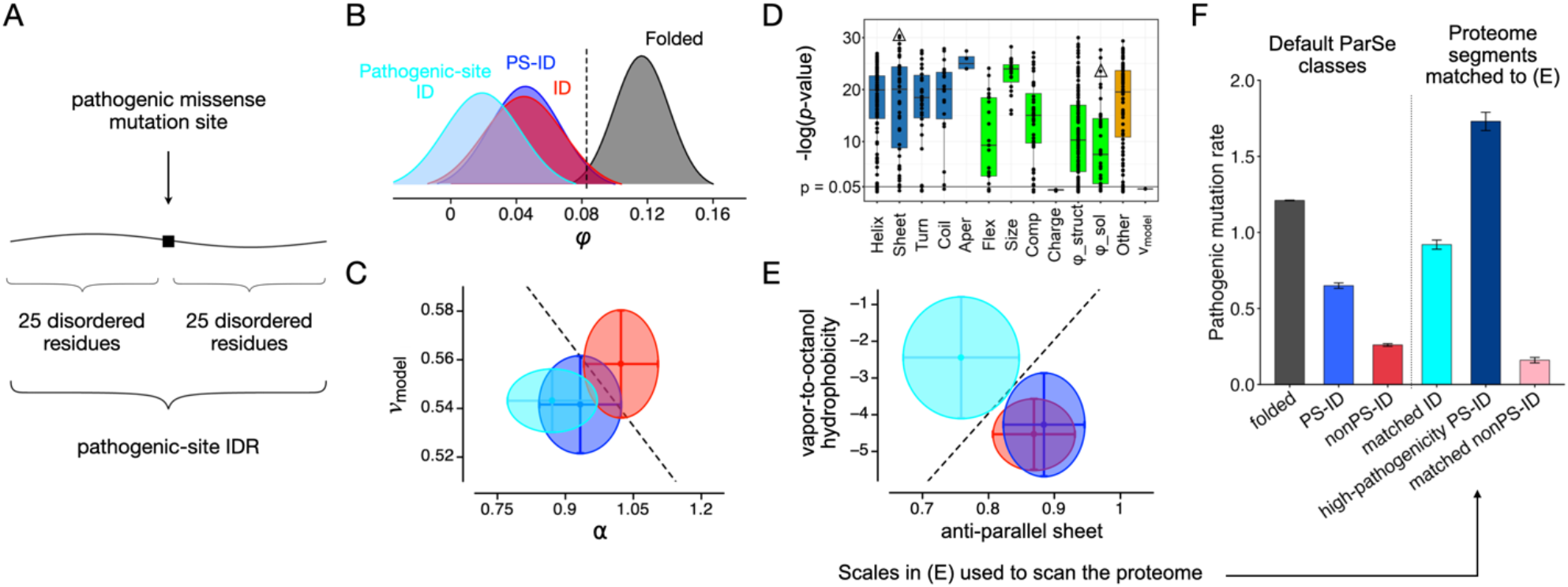
Isolation and recovery of a high-pathogenicity PS-IDR subclass. (**A**) A pathogenic missense mutation flanked by 25 disordered residues on each side define a pathogenic-site IDR. (**B**) Hydrophobicity (*φ*) distributions for pathogenic-site IDRs (cyan) and the PS-ID (blue), conventional ID (red), and folded training sets (black). The dashed line is the default folded-versus-ID decision boundary. (**C**) α-helix propensity versus *ν*_model_ for the same sets, excluding folded. Ellipses are mean ± standard deviation. The dashed line is the default P-versus-D decision boundary. (**D**) Welch’s unequal-variance *t*-test comparing pathogenic-site IDRs with the PS-ID training set across 567 amino-acid scales and *ν*_model_. Values are −log(*p*-value). Scales are grouped by property class. Conformation-based scales (blue boxplots) are α-helix (Helix), sheet or strand (Sheet), β-turn, tight turn, or reverse turn (Turn), coil or loop (Coil), and aperiodic (Aper) propensities. Physicochemical-based scales (green boxplots) are flexibility (Flex), size (Size), composition (Comp), negative charge, positive charge, or net charge (Charge), and hydrophobicity separated into structure-(*φ*_struct) and solution-based (*φ*_sol) scales. Properties not easily mapping into a conformation- or physicochemical-based group were combined separately (Other). Boxplots span the 25^th^ to 75^th^ percentiles; the central bar is the median. Triangles mark the two scales used in (E) that map to orthogonal dimensions of variance in the human proteome. (**E**) Anti-parallel sheet propensity versus vapor-to-octanol hydrophobicity for pathogenic-site IDRs (cyan) and the PS-ID (blue) and conventional ID (red) training sets. Ellipses are mean ± standard deviation. (B-E) compare the pathogenic-site IDR set with the ParSe training sets. (**F**) Normalized pathogenic missense mutation rates in the reference human proteome. The first three bars are default ParSe classes. The next three bars are disordered segments whose sheet and hydrophobicity values match the pathogenic-site IDR centroid in (E). Matched ID is that set of 9,072 segments. High-pathogenicity PS-ID is the α/*ν*_model_ PS split of the matched set. Matched nonPS-ID is the remainder. Error bars are the standard error.

We next identified two orthogonal scales that recover the high-pathogenicity subclass. To find sequence properties that distinguish pathogenic-site IDRs from ordinary PS-IDRs, we screened 566 amino-acid property scales from the Amino Acid Index Database (42) together with a phase-separation hydrophobicity scale (43) and *ν*_model_ (Figure 3). For each scale, we computed the mean value across the 1,026 pathogenic-site IDRs and the 224 PS-ID training set sequences. Welch’s unequal-variance *t*-test (44) indicated that 87% of the scales produce significantly different means between pathogenic-site IDRs and the PS-ID training set (one-tailed *p*-value < 0.05). The nonparametric Mann-Whitney U-test (45) yielded essentially the same result (Figure S4). When scales were grouped by biophysical category (helix propensity, turn propensity, hydrophobicity, flexibility, etc.), every category except charge contained multiple highly significant scales. The single best discriminator was an anti-parallel β-sheet propensity scale (*p*-value = 4.6 × 10^−31^) (46).

Principal component analysis (PCA) of the highest performing scales indicated that approximately 70% of the variance observed across the human proteome is captured by the first two dimensions (59.7% and 12.7%, respectively) (Figure S4). Many of the high-performing scales are therefore correlated, and two dimensions are sufficient to represent most of the property variation. The anti-parallel β-sheet scale aligned closely with the first principal component, while a vapor-to-octanol hydrophobicity scale (*p*-value = 2.9 × 10^−24^) (47) aligned with the second. In the resulting two-dimensional property space, the centroids of the pathogenic-site IDR and PS-ID sets are clearly separated, whereas the conventional-ID training set centroid overlays the PS-ID centroid (Figure 3).

To confirm the effective identification of regions containing pathogenic mutations, a classifier based on these two orthogonal scales was then used to scan the human proteome for disordered segments whose properties match the pathogenic-site IDRs. The search returned 9,072 segments of ≥20 residues. The normalized pathogenic missense mutation rate in this matched set is 0.92 ± 0.03; higher than the default PS-IDR rate (0.65 ± 0.018) yet lower than the rate in folded domains (1.21 ± 0.002). Splitting the same segments with the original α-helix-propensity and *ν*_model_ criteria separates a high-rate PS subclass from a low-rate nonPS subclass, 1.73 ± 0.06 versus 0.16 ± 0.018. We refer to the high-rate subclass as high-pathogenicity PS-IDRs. An independent route through property space thus recovers the same conclusion as boundary shifting. The elevated pathogenic rate belongs to a PS-IDR subclass, not to matched disordered sequence in general.

### High-pathogenicity PS-IDRs are defined by a refined sticker-spacer grammar

We next examined amino-acid composition and local sequence patterning in the high-pathogenicity PS-IDR set defined above, so that low-rate nonPS sequence is not mixed into the counts. The high-pathogenicity PS-IDR set contains 3,909 segments and 164,880 residues. As a comparison group we used conventional nonPS-IDRs, defined as all ParSe-predicted D-labeled domains of ≥20 residues (Figure 4).

**Figure 4.**
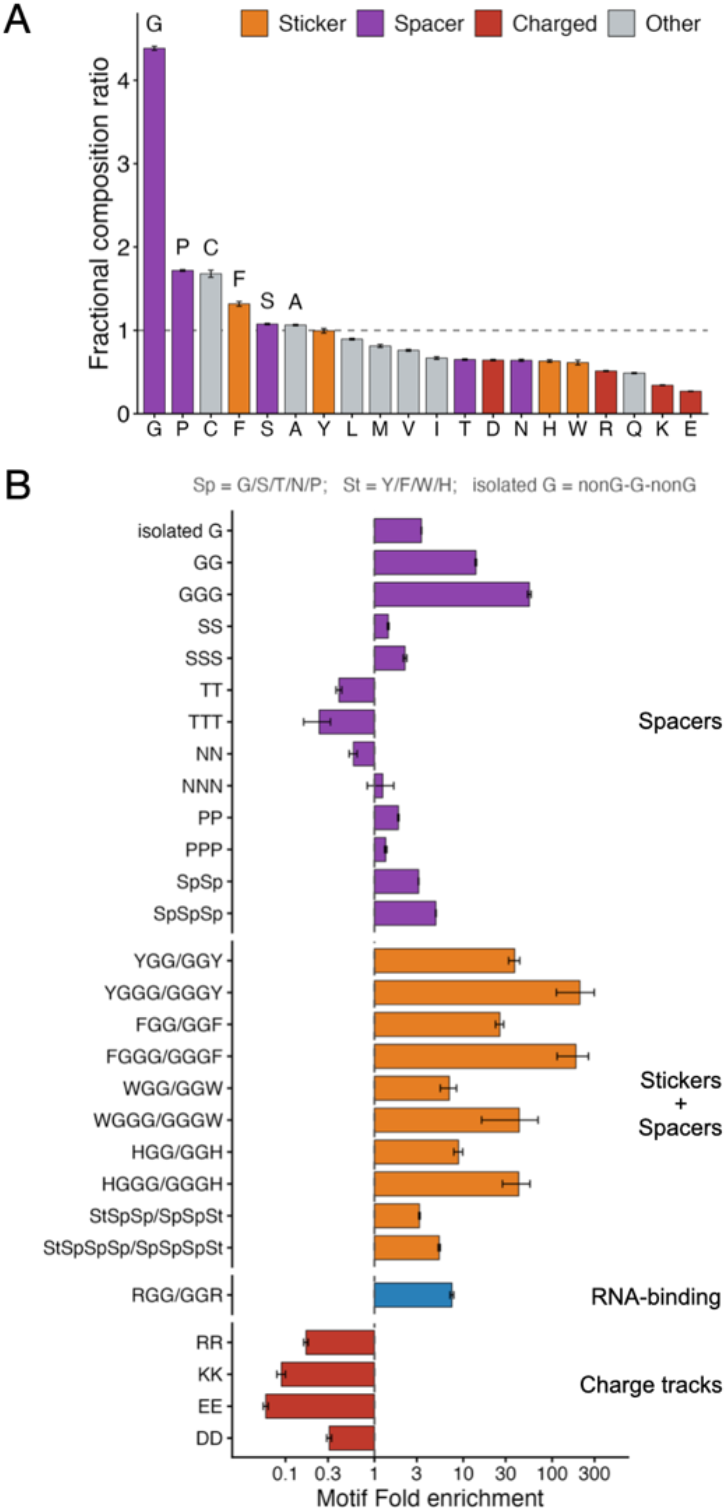
Compositional and motif-level differences between high-pathogenicity PS-IDRs and conventional nonPS-IDRs. (**A**) Ratio of fractional composition of each amino acid in high-pathogenicity PS-IDRs relative to conventional nonPS-IDRs. Amino acids are ordered by enrichment. Bars are colored by functional class: stickers (orange), spacers (purple), charged residues (red), and others (gray). The dashed line indicates equal composition (ratio = 1). (**B**) Fold enrichment of selected sequence motifs in high-pathogenicity PS-IDRs versus conventional nonPS-IDRs (log scale). Motifs are grouped by category: spacers, stickers + spacers, the RNA-binding motif RGG, and charge tracks. Sp = G, S, T, N or P; St = Y, F, W or H; isolated G = nonG-G-nonG. Isolated G is shown for comparison with G-runs. The dashed line indicates no enrichment (fold = 1).

High-pathogenicity PS-IDRs are strongly enriched in the canonical stickers and spacers of phase separation. In the sticker-spacer framework (23, 24), the residues G, S, T, N, and to a lesser extent P function as common spacers that promote chain flexibility, extension, and solubility (48–50). In contrast, Y, F, and W, and to a lesser extent H, act as stickers that mediate the multivalent cohesive interactions that drive condensation, e.g., π–π, cation–π, and hydrophobic contacts (35, 49–53). Consistent with this framework, the spacers G, P, and S, and the sticker F, show the largest increases in high-pathogenicity PS-IDRs relative to conventional nonPS-IDRs. The two sets have near-equivalent Y content. Charge is markedly reduced, especially for E and K. Also depleted is R, though less severely than E or K, while D shows only a mild reduction. This pattern is consistent with reduced overall charge density while still permitting some R-mediated cation–π interactions. Also enriched is C, possibly reflecting additional functional constraints such as redox sensitivity.

Importantly, the differences extend well beyond bulk composition. High-pathogenicity PS-IDRs display a highly specific local sequence grammar in which aromatic stickers are preferentially embedded within short runs of spacers, primarily G-runs (Figure 4, Table S1). Motifs of the form YGG/GGY, YGGG/GGGY, FGG/GGF, and FGGG/GGGF are enriched 30-to >200-fold relative to conventional nonPS-IDRs. Analogous patterns involving W or H are also elevated (7-to >40-fold), as are pure G-runs (GGG is enriched >50-fold). Isolated G (nonG-G-nonG) is enriched only 3.4-fold, lower than the 4.4-fold enrichment of G as an amino acid, so the extra G is not by itself the grammar. The RGG/GGR motif, a hallmark of many experimentally validated phase-separating RNA-binding proteins (54, 55), is enriched (7-fold). In contrast, homotypic charge tracks (RR, KK, EE, DD) are strongly depleted. These patterned motifs are far more discriminatory than composition alone, indicating that the precise arrangement of stickers and spacers, rather than the simple presence of particular amino acids, tracks with the elevated pathogenic enrichment.

Collectively, these results define a refined sticker-spacer grammar that characterizes the disordered sequences most vulnerable to pathogenic mutation. Aromatic residues are preferentially embedded within G-rich (and other spacer-rich) contexts, while homotypic charge tracts are strongly depleted. This specific local patterning, rather than bulk composition alone, appears to underlie the elevated mutational sensitivity of high-pathogenicity PS-IDRs.

### Isolated G, not the enriched motifs, absorb pathogenic substitutions

Within the high-pathogenicity PS-IDR subclass, pathogenic missense mutations outnumber benign ones (822 pathogenic versus 588 benign). In contrast, conventional nonPS-IDRs contain far more benign than pathogenic mutations (892 pathogenic versus 4,236 benign). This striking inversion in the pathogenic-to-benign ratio further shows that disease-associated missense variation is strongly concentrated within a small and compositionally selective PS-ID class.

To understand how pathogenic missense mutations disrupt high-pathogenicity PS-IDR sequences, we compared the spectrum of pathogenic versus benign substitutions within these regions. Notably, 86% of pathogenic missense mutations (704/822) substitute a wild-type G, while only 18% (104/588) of benign mutations occur at G (Figure 5, Table S2). This large difference indicates that G substitutions are poorly tolerated in high-pathogenicity PS-IDRs. Because these regions are extremely G-rich (Figure 4), G presents a large mutational target. The strong enrichment of pathogenic over benign G mutations shows many of these substitutions are deleterious.

**Figure 5.**
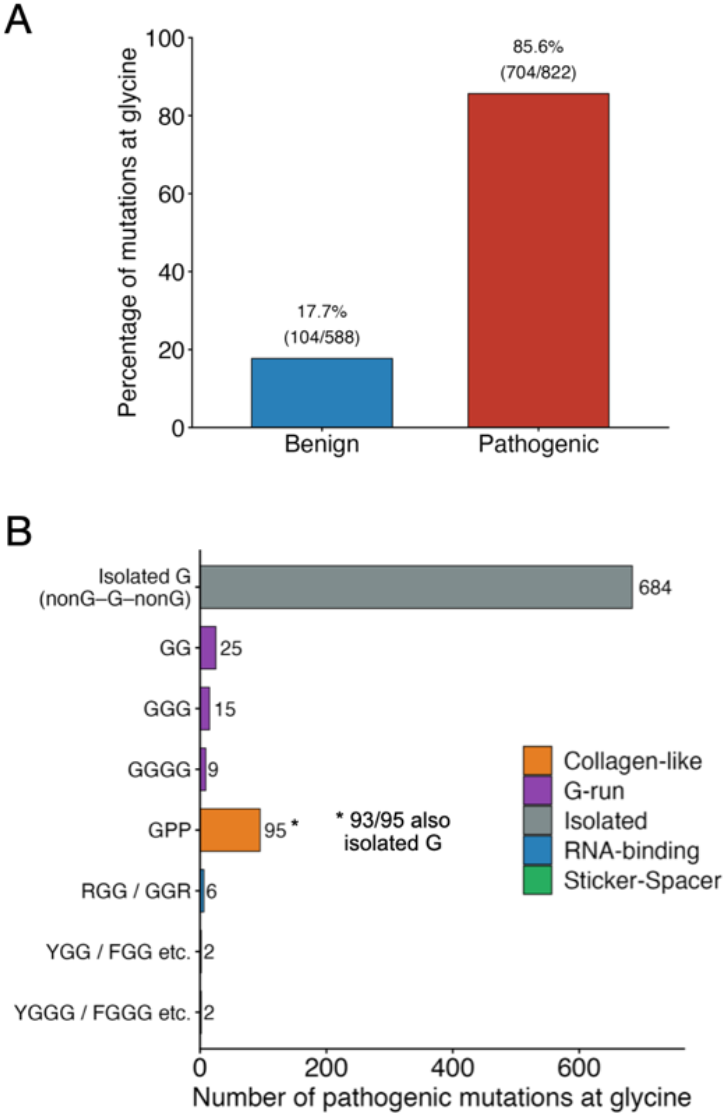
Pathogenic missense mutations in high-pathogenicity PS-IDRs fall mainly at isolated G. (**A**) Percentage of benign versus pathogenic missense mutations that substitute a wild-type G. (**B**) Sequence context of the 704 pathogenic G substitutions. Isolated G is nonG-G-nonG. G-run bars are substitutions at G inside GG, GGG, or GGGG. RGG/GGR and YGG/FGG-type bars are substitutions at G inside those motifs. The GPP bar is the collagen-like motif (95 substitutions at G). Of those 95, 93 are also isolated G and are already included in the isolated-G bar. GPP is shown to mark that subset, not as an added 95 events. Per-occurrence rates are in Table S3.

The overwhelming majority of pathogenic G mutations (684 of 704, 97%) occur in isolated contexts (nonG-G-nonG) rather than within contiguous G-runs. Mutations that fall inside GG, GGG, or GGGG motifs are comparatively rare. This is the opposite of what might be expected if the long G-runs themselves were the primary functional elements being disrupted. Instead, the data suggest that isolated G are more frequently mutated in disease, while the contiguous G-runs are rarely mutated.

Even more striking, the highly enriched sticker-spacer motifs identified earlier (YGG/YGGG, FGG/FGGG, and related patterns) are almost never mutated by either pathogenic or benign variants (Figure 5, Table S2). The RGG motif is likewise rarely hit. Isolated G therefore constitute the primary mutational target. The most straight-forward interpretation is that these patterned motifs are functionally important enough that both pathogenic and benign variation are suppressed.

A notable exception occurs at GPP motifs, which are common in collagen-like sequences (56). Nearly all pathogenic mutations within GPP (95 of 97) substitute the G, whereas most benign mutations at GPP (17 of 19) substitute a P. This pattern matches the known structural requirement for G at every third position in the collagen triple helix (57, 58) and suggests that a subset of sequences classified as high-pathogenicity PS-IDRs contains short collagen-like elements that contribute to the observed G mutational burden.

Raw counts do not correct for how often each context occurs. Isolated G is common in this subclass (163 per 1,000 residues; 26,846 occurrences), whereas YGG/FGG-type motifs are rare (Table S1). Table S3 reports pathogenic and benign substitutions of G per 1,000 motif occurrences. Isolated G remains the main pathogenic target after that correction, 25.5 pathogenic versus 2.3 benign hits per 1,000 occurrences. G-runs and the enriched sticker-spacer motifs stay quiet on a per-occurrence basis, and several have benign density as high as or higher than pathogenic density. GPP has a higher pathogenic density per occurrence (47.2 pathogenic versus 1.0 benign hits per 1,000 occurrences), but 93 of the 95 pathogenic GPP substitutions at G are also isolated G. GPP marks a collagen-like subset of isolated G.

The chemical identity of the substituting residue is less discriminatory (Table S2). A substantial fraction of G substitutions introduce a charged residue (55% of pathogenic and 44% of benign G mutations). This preference may partly reflect codon structure, as single-nucleotide changes in G codons readily generate R, D, or E. However, the similar frequency in both pathogenic and benign sets indicates that the identity of the substituting residue is not a strong determinant of pathogenicity. Common substitutions such as G→S, G→A, G→C, and G→V are also frequent in both sets. Thus, once a G is mutated, the nature of the new residue does not strongly distinguish pathogenic from benign outcomes.

Together, these results both support and refine the earlier conclusions. The extreme rarity of mutations within the most enriched sticker-spacer motifs is consistent with strong functional constraint on local sequence patterning. At the same time, the dominant role of isolated G substitutions, together with the clear collagen-like signature at GPP motifs, indicates that the pathogenic mutational burden arises from more than one source: disruption of phase-separation grammar and, in a subset of cases, violation of collagen-like structural constraints. The high-pathogenicity PS-IDR set therefore appears heterogeneous, containing both classic phase-separating sequences that follow a refined sticker-spacer architecture and sequences that carry additional G-specific structural requirements.

### High-pathogenicity PS-IDRs occur predominantly in regulatory and RNA-related proteins

We identified 2,659 human proteins containing at least one high-pathogenicity PS-IDR of ≥20 residues. To place this subclass in a broader biological context, we examined these proteins using PANTHER Gene Ontology (GO) annotation (Figure 6, Table S4) (59).

**Figure 6.**
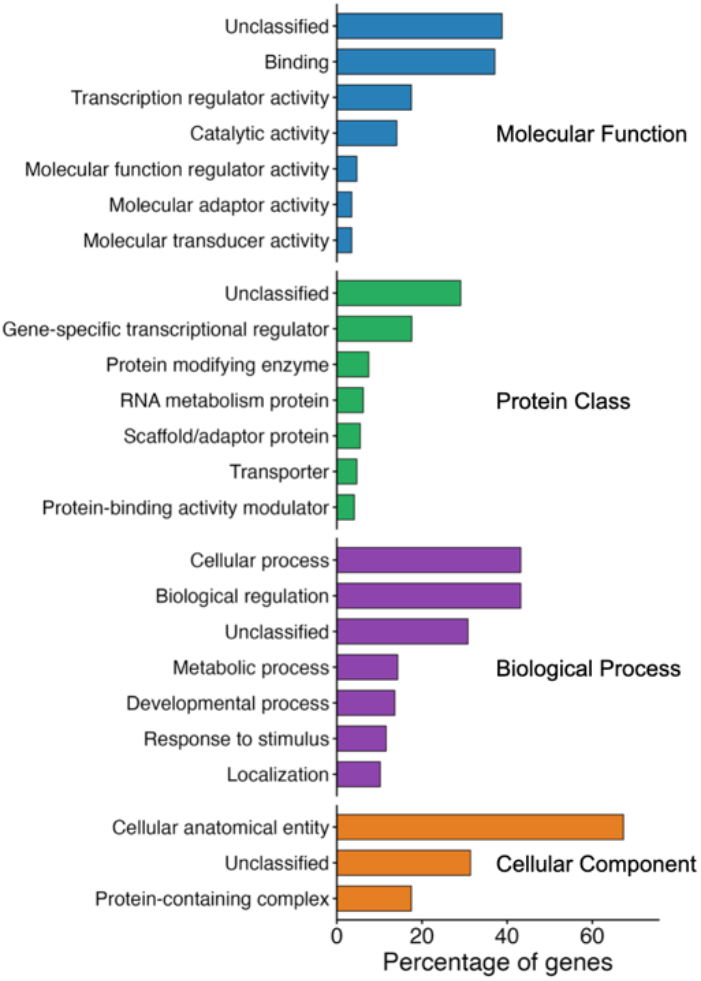
Gene Ontology annotation of proteins containing high-pathogenicity PS-IDRs. PANTHER classification of the 2,659 human proteins that contain at least one high-pathogenicity PS-IDR of ≥20 residues. Bars show the percentage of genes annotated to each term in Molecular Function, Protein Class, Biological Process, and Cellular Component. Terms are ordered within each ontology by that percentage. Unclassified genes are included. Lower-ranked terms are omitted for clarity.

The strongest annotated Molecular Function terms are binding (37.1% of genes) and transcription regulator activity (17.5%), followed by catalytic activity (14.1%) and molecular function regulator activity (4.7%). In the Protein Class ontology, the leading classified category is gene-specific transcriptional regulator (17.6%), with additional representation of protein-modifying enzymes, RNA metabolism proteins, and scaffold/adaptor proteins. These terms match the cellular roles most often associated with phase-separating proteins that use low-complexity or prion-like IDRs to organize transcription, RNA processing, and ribonucleoprotein granules (14, 17, 49). Transcription regulator activity and gene-specific transcriptional regulators correspond to condensate-associated control of gene expression, RNA metabolism proteins to RNA processing and ribonucleoprotein granules, and scaffold/adaptor proteins to the multivalent assemblies that organize those compartments (18, 60, 61). Biological Process annotations are dominated by the broad terms cellular process and biological regulation (each 43.2% of genes). Cellular Component annotations are likewise broad, led by cellular anatomical entity (67.3%) and protein-containing complex (17.5%).

Categories linked to the collagen-like sequence features identified earlier are only modestly represented (Table S4). Extracellular-matrix proteins constitute 2.4% of the set, and structural proteins 1.1%. Thus, while short collagen-like motifs (particularly GPP) contribute to the G-centered mutational spectrum, full-length collagen or collagen-related proteins do not dominate the high-pathogenicity PS-IDR set.

A substantial fraction of proteins remain unclassified in every ontology (29-39% of genes). This is expected for a large set of IDR-rich proteins; many still lack detailed experimental characterization, and GO annotations remain biased toward well-studied proteins and toward functions that can be assigned from structured domains (62, 63). Proteins whose principal functional elements are disordered are therefore often associated with only generic or no GO terms.

Overall, the Gene Ontology profile indicates that high-pathogenicity PS-IDRs occur predominantly in proteins annotated to binding and, more specifically, to regulatory, RNA-related, and scaffolding functions. These are the same biological contexts in which the refined sticker-spacer grammar is expected to operate. Many of the best-characterized phase-separating proteins that rely on aromatic residues embedded in G-rich motifs play critical cellular roles in RNA-centric processes like transcription, RNA processing, and the dynamic assembly of ribonucleoprotein granules or signaling hubs (17, 18, 49). In these settings, weak, multivalent, and reversible interactions are functionally advantageous, consistent with the sequence features enriched in the high-pathogenicity PS-IDR subclass. The collagen-like contribution remains secondary at the level of whole-protein annotation.

## DISCUSSION

Missense variation in disordered protein regions is confounding because most IDRs are comparatively tolerant of substitution (3, 64). This study isolates a core subset of PS-IDRs in which that statement is not true. By using pathogenic mutation rates as an independent biological readout, we demonstrate that mutational vulnerability is concentrated in a narrow compositional and biophysical window. Sequences in this subset are not defined by a generic disordered composition. Rather, they follow a local sticker-spacer architecture with aromatic residues embedded in short G-rich motifs and homotypic charge tracts avoided. The most enriched of those motifs are almost never mutated by either pathogenic or benign substitutions. The main pathogenic target is instead isolated G, a glycine flanked on both sides by non-glycine residues.

That contrast between almost invariant motifs and mutable isolated G explains why IDR variants have been hard to interpret. Averaged over all disordered sequence, missense changes look tolerated. Inside this subclass, a substitution is informative only if its position is specified. A change that breaks a YGG, FGG, or related motif is rare. A change at isolated G is the recurrent pathogenic event, consistent with loss of a single flexible spacer that is not buffered by neighboring G. Biophysical studies on G in condensate-forming proteins support that reading. G keeps droplets fluid and prevents stickers from locking into solid-like states (49, 65, 66).

The proteins that carry these high-pathogenicity segments are enriched in regulatory, RNA-related, and scaffolding roles, matching the cellular contexts in which classic phase-separating proteins operate. The classic sticker-spacer model of phase separation posits that multivalent adhesive motifs (stickers) interspersed with flexible, solubility-promoting elements (spacers) drive the weak, reversible interactions required for condensate formation (23, 24). Our results refine that framework. The sequences with the highest pathogenicity in the human proteome follow a precise local grammar, and it is the local code used in the cellular contexts where phase separation is already known to matter.

The data impose two key limitations on using this framework to interpret missense variants. First, the high-pathogenicity set is heterogeneous. GPP is enriched, and almost all pathogenic GPP substitutions at G are also isolated G. That collagen-like signature is only modestly reflected in whole-protein GO terms. Variant interpretation should therefore ask whether an isolated G sits in GPP, not only whether the region is called a PS-IDR. Second, bulk amino-acid composition is a weak discriminator. Composition ratios are modest next to motif enrichments of 7-to >200-fold. Predictors or annotations that stop at residue frequencies will miss the sites that matter.

In practical terms, a missense variant in an IDR should be scored in three steps: whether it lies in the high-pathogenicity PS-IDR subclass, whether it falls in a sticker-spacer motif, at G in GPP, or at isolated G, and whether the surrounding protein functions in condensate-associated regulation. A substitution inside YGG, FGG, or a related motif is rare and should be treated as a possible grammar-breaking change. A substitution at G in GPP has a high pathogenic density and a known structural rule. Isolated G carries most of the observed pathogenic burden, but it is also a frequent context, so it is not a standalone predictor. This provides disordered sequence a site-resolved basis that folded-domain stability arguments already have. In this way, the present work advances the sticker-spacer model from a general organizing principle into a context-dependent sequence grammar that is demonstrably linked to mutational vulnerability. This provides both a mechanistic explanation for the elevated mutational vulnerability of these sequences and a practical framework for interpreting genetic variants in disordered protein regions.

## EXPERIMENTAL PROCEDURES

### Protein and genetic variation databases

The one-sequence-per-gene version of the reference human proteome (UP000005640) was downloaded from UniProtKB (39) in May 2026. The human variants index (humsavar.txt, release 2026_01) was downloaded from UniProt in March 2026.

### Sequence-based prediction of domain organization and phase separation potential

Domain-level organization and phase separation (PS) potential were calculated with ParSe v2 (22, 25), available as source code and as a webtool at github.com/stevewhitten.

### Comparison of ParSe v2 with other predictors

ID predictions used for comparison with ParSe v2 were obtained from Metapredict (metapredict.net) and PUNCH2 (alienlabs.ucd.ie/punch2) (29, 30). Phase separation predictions used human proteome scores obtained from the supplemental datasets reported with DeePhase, catGRANULE2, FuzDrop, PSPHunter, PhaSePred, PSAP, and PSPire (31–34, 36– 38). PSCORE predictions were calculated with the Python implementation and associated database files available at doi.org/10.7554/eLife.31486.022 (35).

### Normalized missense mutation rate

Pathogenic rates were calculated from missense variants labeled pathogenic or likely pathogenic. Benign rates were calculated from missense variants labeled benign or likely benign. For a given region class, the unnormalized mutation proportion is,

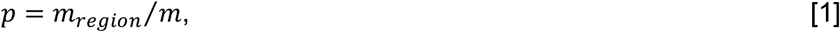

where *m*_*region*_ is the number of mutations in that class and *m* is the total number of mutations of the same clinical class in the sequence set. The normalized rate is,

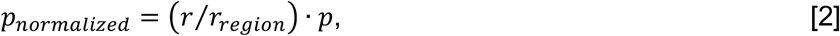

where *r*_*region*_ is the number of residues in that class and *r* is the total number of residues in the sequence set. This is equivalent to dividing the mutation proportion by the residue proportion. The standard error of *p* is,

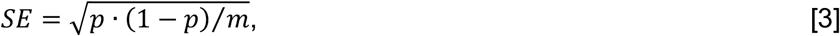

and the standard error of the normalized rate is,

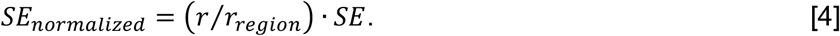

### Motif fold enrichment

Motif frequency was expressed per 1000 residues,

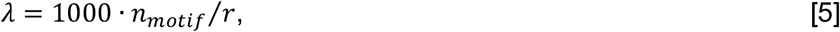

where *n*_*motif*_ is the number of motif occurrences in a sequence set. The standard error is,

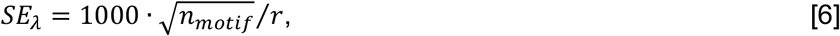

Fold enrichment of a motif in set A relative to set B is,

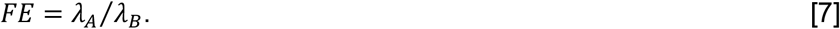

with propagated standard error,

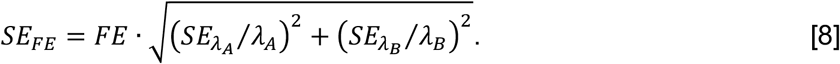

### Statistical tests and principal component analysis

One-tailed *p*-values for differences in mean sequence property values were calculated with Welch’s unequal-variance *t*-test (44) and the Mann-Whitney U test (45) in the statistical program R (67). Rank concordance between ParSe v2 PS potential and other proteome-wide phase-separation scores was measured with Spearman’s *ρ*in R. Principal component analysis was performed in R after centering each variable at zero and scaling to unit variance.

### Gene Ontology classification

Protein classification used the PANTHER classification system (59) at www.pantherdb.org accessed in August 2026.

## Data availability

Source code for calculating domain-level organization, including high-pathogenicity PS-IDRs, is available at github.com/stevewhitten as Fortran, Python, and browser-based HTML implementations. The folded, conventional ID, and PS-ID training sets used for ParSe v2, and the pathogenic-site IDR set used here, are available at the same site.

## Supporting information

This article contains supporting information (29–39, 46, 47, 59).

### Funding and additional information

This work was supported by the National Science Foundation under grant 1943488 (to L. E. H.), and Texas State University Office of Research and Sponsored Projects through the Research Enhancement Program (to K. A. L. and S. T. W.).

The content is solely the responsibility of the authors and does not necessarily represent the official views of the National Science Foundation.

## Abbreviations

The abbreviations used are

ID: intrinsic disorder
IDR: intrinsically disordered region
PS-ID: phase-separating intrinsic disorder
PS-IDR: phase-separating intrinsically disordered region
v2: version 2
SE: standard error.

