## Supporting Information for "A refined sticker-spacer grammar marks disordered regions with interpretable pathogenic missense variation"

**Contents:**

Supporting Tables
S1. Fold enrichment of selected sequence motifs in high-pathogenicity PS-IDRs versus conventional nonPS-IDRs.
S2. Pathogenic and benign missense mutations within high-pathogenicity PS-IDRs.
S3. Pathogenic and benign missense density at glycine in high-pathogenicity PS-IDRs.
S4. PANTHER annotation of 2,659 human proteins that contain at least one high-pathogenicity PS-IDR of ≥20 residues.

Supporting Figures
S1. Disorder predictors recover the ParSe folded-versus-ID split but do not separate PS-IDRs from conventional IDRs.
S2. ParSe v2 agrees in rank with most sequence-based phase-separation predictors, but not with structure-aware or no-IDR-targeted methods.
S3. Benign missense mutation rates are insensitive to ParSe v2 boundary shifts.
S4. Property scales that distinguish pathogenic-site IDRs from the PS-ID training set, and the principal axes of proteome variation.

Supporting References

**Table S1. Fold enrichment of selected sequence motifs in high-pathogenicity PS-IDRs versus conventional nonPS-IDRs.** Frequencies are motif counts per 1,000 residues. Fold enrichment is the high-pathogenicity PS-IDR frequency divided by the conventional nonPS-IDR frequency. Standard errors were calculated as described in Experimental Procedures. Isolated G are nonG-G-nonG.

| | High-pathogenicity PS-IDRs | | nonPS-IDRs | | | |
|---|---|---|---|---|---|---|
| **Motif** | **Frequency per 1000 residues** | **Standard Error** | **Frequency per 1000 residues** | **Standard Error** | **Fold Enrichment** | **Standard Error** |
| **Spacers, Sp = G, S, T, N, or P** | | | | | | |
| Isolated G | 162.82 | 0.99 | 48.12 | 0.20 | 3.38 | 0.02 |
| GG | 56.64 | 0.59 | 4.08 | 0.06 | 13.88 | 0.25 |
| GGG | 23.68 | 0.38 | 0.43 | 0.02 | 55.45 | 2.63 |
| SS | 15.11 | 0.30 | 10.54 | 0.09 | 1.43 | 0.03 |
| SSS | 3.68 | 0.15 | 1.66 | 0.04 | 2.21 | 0.10 |
| TT | 1.09 | 0.08 | 2.72 | 0.05 | 0.40 | 0.03 |
| TTT | 0.05 | 0.02 | 0.22 | 0.01 | 0.24 | 0.08 |
| NN | 0.53 | 0.06 | 0.92 | 0.03 | 0.58 | 0.06 |
| NNN | 0.07 | 0.02 | 0.05 | 0.01 | 1.24 | 0.41 |
| PP | 34.37 | 0.46 | 18.47 | 0.13 | 1.86 | 0.03 |
| PPP | 7.51 | 0.21 | 5.61 | 0.07 | 1.34 | 0.04 |
| SpSp | 321.90 | 1.40 | 102.62 | 0.30 | 3.14 | 0.02 |
| SpSpSp | 186.27 | 1.06 | 37.84 | 0.18 | 4.92 | 0.04 |
| SpSpSpSp | 115.57 | 0.84 | 14.93 | 0.11 | 7.74 | 0.08 |
| **Stickers + Spacers, St = Y, F, W, or H** | | | | | | |
| YG/GY | 5.30 | 0.18 | 0.89 | 0.03 | 5.93 | 0.27 |
| YGG/GGY | 1.87 | 0.11 | 0.05 | 0.01 | 37.79 | 5.41 |
| YGGG/GGGY | 0.87 | 0.07 | 0.004 | 0.002 | 204.94 | 93.23 |
| FG/GF | 9.61 | 0.24 | 1.31 | 0.03 | 7.32 | 0.26 |
| FGG/GGF | 2.50 | 0.12 | 0.10 | 0.01 | 25.78 | 2.73 |
| FGGG/GGGF | 1.10 | 0.08 | 0.01 | 0.006 | 185.02 | 71.26 |
| WG/GW | 1.27 | 0.09 | 0.62 | 0.02 | 2.06 | 0.16 |
| WGG/GGW | 0.28 | 0.04 | 0.04 | 0.01 | 6.96 | 1.44 |
| WGGG/GGGW | 0.11 | 0.03 | 0.003 | 0.001 | 42.70 | 26.63 |
| HG/GH | 5.65 | 0.19 | 2.08 | 0.04 | 2.71 | 0.10 |
| HGG/GGH | 1.02 | 0.08 | 0.12 | 0.01 | 8.84 | 1.02 |
| HGGG/GGGH | 0.36 | 0.05 | 0.01 | 0.003 | 41.99 | 14.36 |
| StSpSp/SpSpSt | 25.89 | 0.40 | 8.07 | 0.08 | 3.21 | 0.06 |
| StSpSpSp/SpSpSpSt | 14.38 | 0.30 | 2.68 | 0.05 | 5.37 | 0.15 |
| StSpSpSpSp/SpSpSpSpSt | 8.69 | 0.23 | 1.00 | 0.03 | 8.69 | 0.34 |
| **RNA-binding** | | | | | | |
| RGG/GGR | 4.62 | 0.17 | 0.62 | 0.02 | 7.45 | 0.36 |
| **Charge tracks** | | | | | | |
| RR | 1.48 | 0.09 | 8.95 | 0.09 | 0.17 | 0.01 |
| KK | 1.16 | 0.08 | 12.32 | 0.10 | 0.09 | 0.01 |
| EE | 1.69 | 0.10 | 27.61 | 0.15 | 0.06 | 0.004 |
| DD | 1.75 | 0.10 | 5.64 | 0.07 | 0.31 | 0.02 |
| **Negative, Neg = D or E** | | | | | | |

| | | | | | | |
|---|---|---|---|---|---|---|
| NegNeg | 6.82 | 0.20 | 52.34 | 0.21 | 0.13 | 0.004 |
| **Positive, Pos = K or R** | | | | | | |
| PosPos | 4.27 | 0.16 | 35.34 | 0.17 | 0.12 | 0.005 |
| **Adjacent opposite charge** | | | | | | |
| PosNeg/NegPos | 14.79 | 0.30 | 51.61 | 0.21 | 0.29 | 0.01 |
| **Collagen motif** | | | | | | |
| GPP | 12.20 | 0.27 | 0.96 | 0.03 | 12.68 | 0.47 |

**Table S2. Pathogenic and benign missense mutations within high-pathogenicity PS-IDRs.** Counts are from the UniProt humsavar index (1). “At the G” and “at a G” gives the subset of motif mutations that substitute a wild-type G. Isolated G are nonG-G-nonG.

| | Pathogenic missense mutations **count** | Benign missense mutations **count** |
|---|---|---|
| Total within high-pathogenicity PS-IDRs | 822 | 588 |
| **WT residue is G:** | 704 | 104 |
| WT residue is G, mutation is R | 175 | 24 |
| WT residue is G, mutation is K | 0 | 0 |
| WT residue is G, mutation is D | 124 | 12 |
| WT residue is G, mutation is E | 78 | 8 |
| WT residue is G, mutation is Y | 0 | 0 |
| WT residue is G, mutation is F | 0 | 0 |
| WT residue is G, mutation is W | 5 | 0 |
| WT residue is G, mutation is H | 0 | 0 |
| WT residue is G, mutation is S | 126 | 30 |
| WT residue is G, mutation is T | 1 | 0 |
| WT residue is G, mutation is N | 0 | 0 |
| WT residue is G, mutation is P | 0 | 0 |
| WT residue is G, mutation is A | 25 | 11 |
| WT residue is G, mutation is C | 64 | 5 |
| WT residue is G, mutation is I | 0 | 0 |
| WT residue is G, mutation is L | 2 | 0 |
| WT residue is G, mutation is V | 104 | 14 |
| WT residue is G, mutation is M | 0 | 0 |
| WT residue is G, mutation is Q | 0 | 0 |
| **Mutation at a WT G, flanked by:** | | |
| nonG-G-nonG | 684 | 62 |
| nonG-GG-nonG | 9 | 22 |
| nonG-GGG-nonG | 6 | 10 |
| nonG-GGGG-nonG | 2 | 4 |
| **Mutation within a motif:** | | |
| GG | 25 | 46 |
| GGG | 15 | 26 |
| GGGG | 9 | 14 |
| GPP (at the G) | 97 (95) | 19 (2) |
| RGG / GGR (at a G) | 10 (6) | 6 (4) |
| YGG / GGY (at a G) | 1 (1) | 2 (2) |
| YGGG / GGGY (at a G) | 2 (2) | 0 (0) |
| FGG / GGF (at a G) | 1 (1) | 4 (4) |
| FGGG / GGGF (at a G) | 0 (0) | 2 (2) |
| WGG / GGW (at a G) | 0 (0) | 0 (0) |
| WGGG / GGGW (at a G) | 0 (0) | 0 (0) |
| HGG / GGH (at a G) | 0 (0) | 2 (0) |
| HGGG / GGGH (at a G) | 0 (0) | 2 (0) |

**Table S3. Pathogenic and benign missense density at glycine in high-pathogenicity PS-IDRs.** Counts are substitutions of a wild-type G only, from Table S2. Isolated G is nonG-G-nonG. Occurrences use the 164,880-residue high-pathogenicity PS-IDR set and the motif frequencies in Table S1. Hits are per 1,000 occurrences = 1,000 $\times$ count / occurrences. Standard errors were calculated as described for frequencies in Experimental Procedures. Of 95 pathogenic GPP substitutions at G, 93 are also isolated G; GPP is not an independent class in this table. A dash means the pathogenic/benign ratio is undefined. Total mutations in the subclass: 822 pathogenic, 588 benign.

| Motif | Frequency per 1,000 residues | Occurrences | Pathogenic (at G) | Pathogenic hits per 1,000 occurrences | Benign (at G) | Benign hits per 1,000 occurrences | Pathogenic/Benign |
|---|---|---|---|---|---|---|---|
| Isolated G | 162.82 ± 0.99 | 26,846 | 684 | 25.48 ± 0.97 | 62 | 2.31 ± 0.29 | 11.03 ± 1.46 |
| GPP | 12.20 ± 0.27 | 2,012 | 95 | 47.22 ± 4.84 | 2 | 0.99 ± 0.70 | 47.50 ± 33.94 |
| RGG / GGR | 4.62 ± 0.17 | 761 | 6 | 7.88 ± 3.22 | 4 | 5.26 ± 2.63 | 1.50 ± 0.97 |
| YGG / GGY | 1.87 ± 0.11 | 308 | 1 | 3.25 ± 3.25 | 2 | 6.49 ± 4.59 | 0.50 ± 0.61 |
| YGGG / GGGY | 0.87 ± 0.07 | 144 | 2 | 13.89 ± 9.82 | 0 | 0 | - |
| FGG / GGF | 2.50 ± 0.12 | 413 | 1 | 2.42 ± 2.42 | 4 | 9.69 ± 4.84 | 0.25 ± 0.28 |
| FGGG / GGGF | 1.10 ± 0.08 | 182 | 0 | 0 | 2 | 10.99 ± 7.77 | 0 |
| WGG / GGW | 0.28 ± 0.04 | 46 | 0 | 0 | 0 | 0 | - |
| WGGG / GGGW | 0.11 ± 0.03 | 18 | 0 | 0 | 0 | 0 | - |
| HGG / GGH | 1.02 ± 0.08 | 169 | 0 | 0 | 0 | 0 | - |
| HGGG / GGGH | 0.36 ± 0.05 | 59 | 0 | 0 | 0 | 0 | - |
| GG | 56.64 ± 0.59 | 9,339 | 25 | 2.68 ± 0.54 | 46 | 4.93 ± 0.73 | 0.54 ± 0.14 |
| GGG | 23.68 ± 0.38 | 3,904 | 15 | 3.84 ± 0.99 | 26 | 6.66 ± 1.31 | 0.58 ± 0.19 |

**Table S3. PANTHER annotation of 2,659 human proteins that contain at least one high-pathogenicity PS-IDR of ≥20 residues.** For each term: number of annotated genes, percent of the 2,659 proteins annotated to that term, and percent of all class hits assigned to that term (2). A protein may receive more than one term, so gene-based percentages can sum to more than 100%. “No PANTHER category is assigned” is the unclassified fraction.

| GO term name | number of genes annotated to that term | percent of genes annotated to that term | percent of function hits assigned to that term |
|---|---|---|---|
| **Molecular Function** | | | |
| transporter activity | 108 | 4.20% | 3.30% |
| No PANTHER category is assigned | 994 | 38.80% | 30.10% |
| translation regulator activity | 9 | 0.40% | 0.30% |
| cargo receptor activity | 3 | 0.10% | 0.10% |
| transcription regulator activity | 447 | 17.50% | 13.60% |
| catalytic activity (GO:0003824) | 362 | 14.10% | 11.00% |
| molecular function regulator activity | 120 | 4.70% | 3.60% |
| cytoskeletal motor activity | 10 | 0.40% | 0.30% |
| ATP-dependent activity | 41 | 1.60% | 1.20% |
| molecular transducer activity | 89 | 3.50% | 2.70% |
| molecular adaptor activity | 90 | 3.50% | 2.70% |
| antioxidant activity | 2 | 0.10% | 0.10% |
| structural molecule activity | 73 | 2.90% | 2.20% |
| binding | 950 | 37.10% | 28.80% |
| **Protein Class** | | | |
| extracellular matrix protein | 61 | 2.40% | 2.40% |
| cytoskeletal protein | 98 | 3.80% | 3.80% |
| transporter | 121 | 4.70% | 4.70% |
| scaffold/adaptor protein | 142 | 5.50% | 5.50% |
| DNA metabolism protein | 28 | 1.10% | 1.10% |
| cell adhesion molecule | 31 | 1.20% | 1.20% |
| intercellular signal molecule | 40 | 1.60% | 1.60% |
| protein-binding activity modulator | 105 | 4.10% | 4.10% |
| viral or transposable element protein | 2 | 0.10% | 0.10% |
| RNA metabolism protein | 158 | 6.20% | 6.20% |
| calcium-binding protein | 4 | 0.20% | 0.20% |
| gene-specific transcriptional regulator | 450 | 17.60% | 17.60% |
| defense/immunity protein | 17 | 0.70% | 0.70% |
| translational protein | 19 | 0.70% | 0.70% |
| metabolite interconversion enzyme | 90 | 3.50% | 3.50% |
| protein modifying enzyme | 192 | 7.50% | 7.50% |
| chromatin/chromatin-binding or -regulatory protein | 65 | 2.50% | 2.50% |
| transfer/carrier protein | 8 | 0.30% | 0.30% |
| membrane traffic protein | 68 | 2.70% | 2.70% |
| chaperone | 14 | 0.50% | 0.50% |
| cell junction protein | 6 | 0.20% | 0.20% |
| structural protein | 27 | 1.10% | 1.10% |
| transmembrane signal receptor | 71 | 2.80% | 2.80% |
| No PANTHER category is assigned | 744 | 29.10% | 29.10% |
| **Biological Process** | | | |
| cellular process | 1107 | 43.20% | 23.90% |
| reproductive process | 20 | 0.80% | 0.40% |
| No PANTHER category is assigned | 788 | 30.80% | 17.00% |
| localization | 260 | 10.20% | 5.60% |

| | | | |
|---|---|---|---|
| biological process involved in interspecies interaction between organisms | 15 | 0.60% | 0.30% |
| reproduction | 20 | 0.80% | 0.40% |
| biological regulation | 1107 | 43.20% | 23.90% |
| response to stimulus | 297 | 11.60% | 6.40% |
| homeostatic process | 19 | 0.70% | 0.40% |
| developmental process | 349 | 13.60% | 7.50% |
| rhythmic process | 6 | 0.20% | 0.10% |
| multicellular organismal process | 252 | 9.80% | 5.40% |
| locomotion | 4 | 0.20% | 0.10% |
| metabolic process | 367 | 14.30% | 7.90% |
| growth | 3 | 0.10% | 0.10% |
| immune system process | 27 | 1.10% | 0.60% |
| **Cellular Component** | | | |
| cellular anatomical entity | 1723 | 67.30% | 57.90% |
| protein-containing complex | 449 | 17.50% | 15.10% |
| No PANTHER category is assigned | 805 | 31.40% | 27.00% |

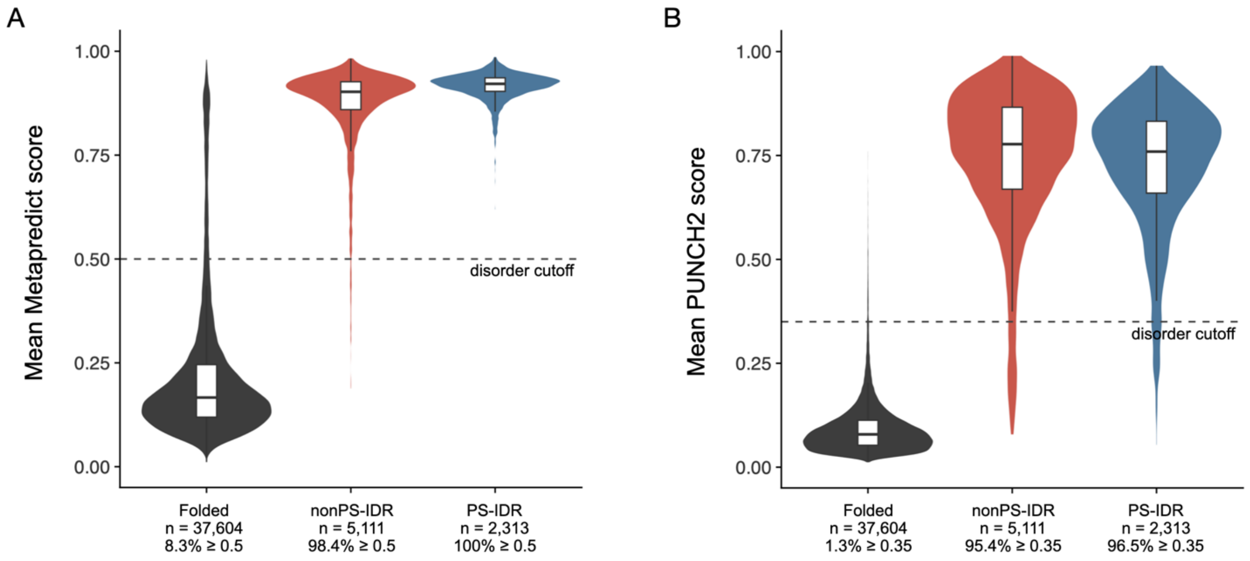


**Figure S1. Disorder predictors recover the ParSe folded-versus-ID split but do not separate PS-IDRs from conventional IDRs.** Mean per-region disorder scores for ParSe v2 segments ≥50 residues from the reference human proteome. (**A**) Metapredict (3); the dashed line is the published disorder cutoff of 0.5. (**B**) PUNCH2 (4); the dashed line is the published disorder cutoff of 0.35. Violin width is scaled within each class; boxes show the median and interquartile range. Labels give the number of regions and the percentage scoring at or above the cutoff.

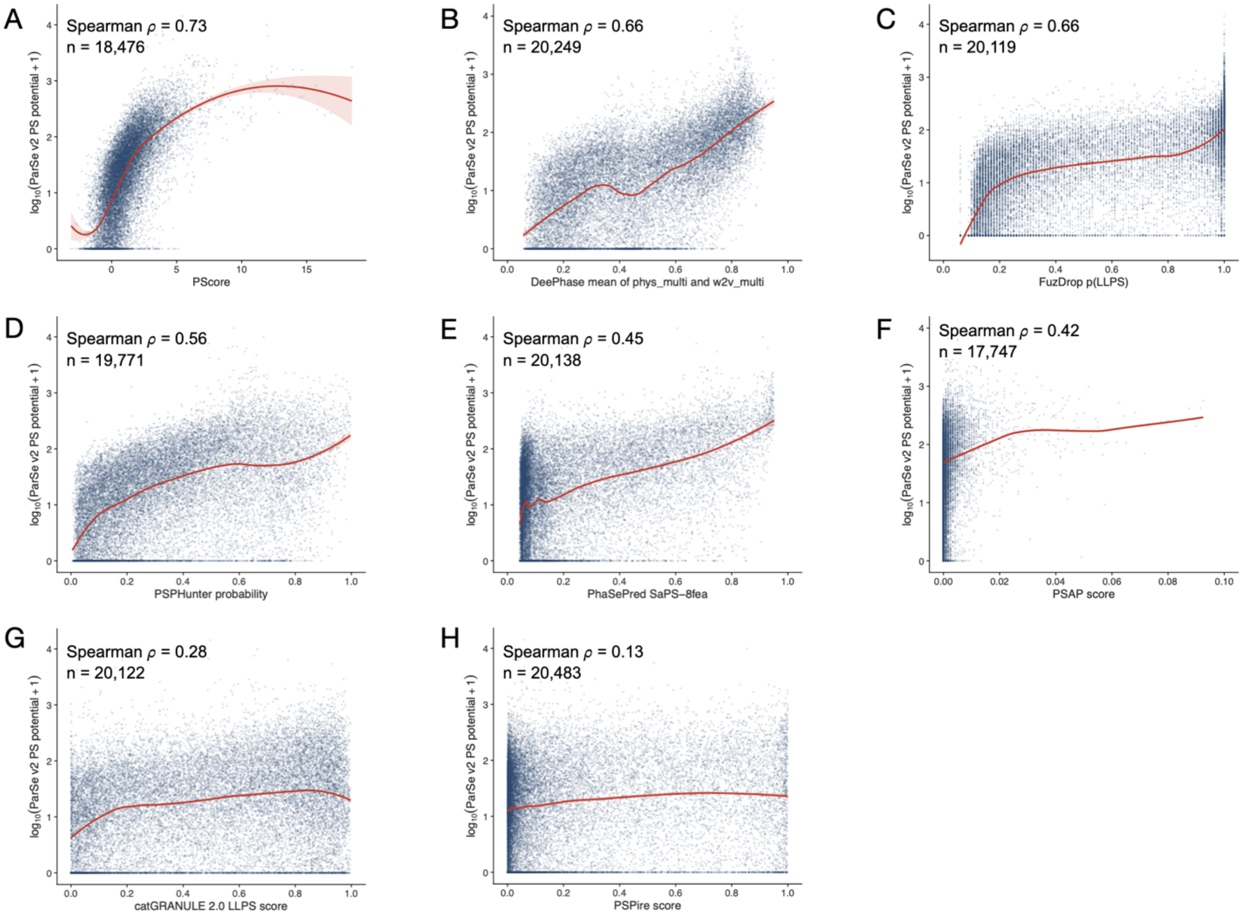


**Figure S2. ParSe v2 agrees in rank with most sequence-based phase-separation predictors, but not with structure-aware or no-IDR-targeted methods.** Pairwise comparison of proteome-wide phase-separation scores with the ParSe v2 PS potential (sum of classifier distances over P-labeled windows). Each point is one human protein. The y-axis is $\log_{10}$(ParSe v2 PS potential + 1) so that the heavy-tailed ParSe scores remain visible. Spearman $\rho$ reports concordance of rank order and is used because the predictors have incompatible numeric ranges and nonlinear score-score relationships. With n≈18,000-20,500 even a very small $\rho$ is statistically nonzero, so the value itself is interpreted as strong (0.6-0.7), moderate (0.4-0.5), or weak (<0.3). (**A**) PScore (5). (**B**) DeePhase mean of phys_multi and w2v_multi (6). (**C**) FuzDrop p(LLPS) (7). (**D**) PSPHunter probability (8). (**E**) PhaSePred SaPS-8fea (9). (**F**) PSAP (10). (**G**) catGRANULE 2.0 LLPS score (11). (**H**) PSPire score (12). PScore, DeePhase, and FuzDrop fall in the strong range. PhaSePred SaPS-8fea and PSAP are moderate. catGRANULE 2.0 and PSPire are weak. Those two exceptions are expected from their design: catGRANULE 2.0 includes AlphaFold-derived structural features and is trained on broadly annotated condensate proteins, and PSPire is built to recover phase-separating proteins that lack long IDRs. ParSe v2 scores only disordered P-labeled windows, so disagreement with those tools is a difference in target class, not a failure of either method. Red lines are LOESS (locally estimated scatterplot smoothing), except in panel F. PSAP scores take only a few discrete values, so the line is fit to the average ParSe score at each of those values.

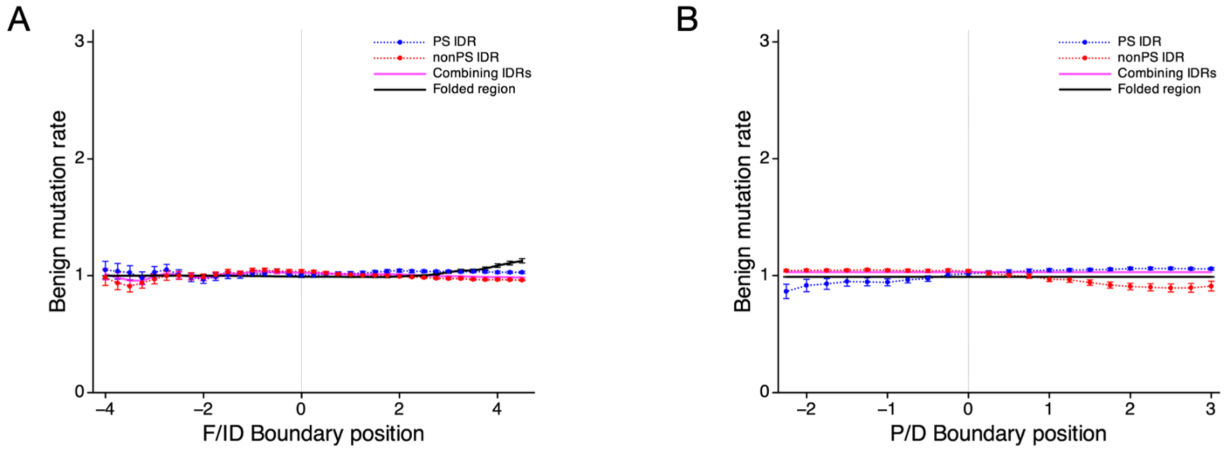


**Figure S3. Benign missense mutation rates are insensitive to ParSe v2 boundary shifts.**
Normalized benign rates for folded regions, PS-IDRs, conventional IDRs, and combined IDRs after shifts of (**A**) the folded-versus-ID boundary and (**B**) the P-versus-D boundary. Error bars are the standard error.

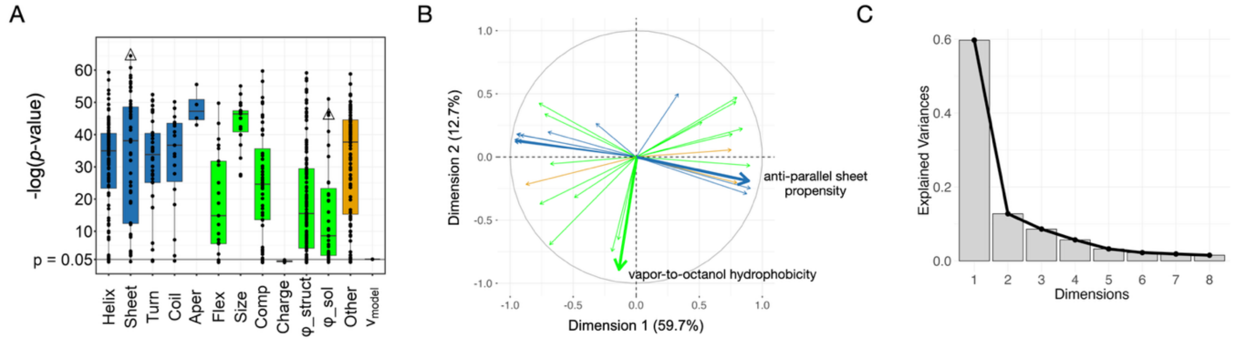


**Figure S4. Property scales that distinguish pathogenic-site IDRs from the PS-ID training set, and the principal axes of proteome variation.** (**A**) Mann-Whitney U test comparing pathogenic-site IDRs with the PS-ID training set across 567 amino-acid scales and $\nu_{\mathrm{model}}$. Values are −log(*p*-value). Scales are grouped by property class. 91% of scales have significantly different means in the two sets (one-tailed *p*-value < 0.05). Boxes span the 25$^{\mathrm{th}}$ to 75$^{\mathrm{th}}$ percentiles; the center line is the median. Triangles mark the two scales labeled in (B). (**B**) PCA biplot of proteome variation using the three lowest *p*-value scales from each property class in (A). Charge-based scales (each with *p*-value > 0.05) and $\nu_{\mathrm{model}}$ (*p*-value ~ 0.05) were omitted. Dimension 1 (59.7%) aligns with an anti-parallel β-sheet propensity (13), the scale with the lowest *p*-value. Dimension 2 (12.7%) aligns with a vapor-to-octanol hydrophobicity scale (14). (**C**) Scree plot of variance explained by each principal component.
